# The identification and characterization of hillocks in the postnatal mouse and human airway

**DOI:** 10.1101/2025.04.15.648833

**Authors:** Yohana Otto, Gergana Shipkovenska, Qiaozhen Liu, Jiawei Sun, Lida P. Hariri, Viral S. Shah, Jayaraj Rajagopal

## Abstract

Adult lung regeneration has been deeply scrutinized over the past decade. In contrast, the injury resistance mechanisms of the neonatal and pediatric airway have received considerably less attention despite the manifest clinical importance. We recently reported the discovery of the airway hillock in adult murine and human airways. Adult hillocks are stratified structures with luminal squamous barrier cells overlying a dedicated basal stem cell population. Functionally, hillocks serve as an injury-resistant reservoir of dedicated hillock stem cells that can resurface and repopulate the airway epithelium after severe damage. Indeed, hillock basal stem cells undergo massive clonal expansion in the process of repopulating denuded airway epithelium. Since the postnatal lung encounters injuries that can result in airway epithelial denudation in the setting of respiratory infection or aspiration, we sought to assess whether hillocks are present in the neonatal airway. In this manuscript we identify and characterize hillocks in postnatal mouse and human pediatric airways. We show that hillocks are present in mice from postnatal day 3 onwards and that they expand in size through adulthood. The earliest hillocks are functionally immature, but they acquire their injury resistance properties over the course of postnatal maturation. By re-analyzing published pediatric scRNAseq data, we identify cells with a hillock squamous cell gene signature that is conserved across species. Finally, we identify bona fide hillocks in an 8-month-old infant and an 8-year-old child. We now wonder whether the presence and maturation of hillocks has implications for disorders of the neonatal and childhood airway. More specifically, we hypothesize that the incomplete maturation of hillocks could create a window of vulnerability in neonates who may be particularly susceptible to airway damage in the setting of infection or aspiration. Thus, there is a need to define when human hillocks first form and establish the time window during which they functionally mature as a prelude to determining whether the appearance and properties of hillocks correlate to clinical phenotypes and outcomes.

---

Adult lung regeneration has been deeply scrutinized in the past decade, however, the injury resistance mechanisms of the neonatal and pediatric airway have been understudied despite their clinical significance. We recently reported the discovery of the airway hillock in adult airway of mouse and man (1, 2). Hillocks are stratified structures with luminal squamous barrier cells overlying a dedicated basal stem cell population. Functionally, hillocks serve as an injury-resistant reservoir of dedicated stem cells that can resurface and repopulate the airway epithelium after severe damage (1). Indeed, hillock basal stem cells undergo massive clonal expansion in the process of repopulating denuded airway epithelium. Since the postnatal lung encounters injuries that can result in airway epithelial denudation in the setting of respiratory infection or aspiration, we sought to assess whether hillocks are present in the neonatal airway. If so, we hypothesized that neonatal and pediatric hillocks might contribute to the resiliency of the young airway as it encounters new environmental threats outside the protective maternal milieu.

We previously defined adult murine hillocks by a series of criteria including keratin 13 (Krt13) expression in squamous luminal cells and underlying hillock basal stem cells, the absence of ciliated cells, and their occurrence in characteristic locations over the posterior membrane and over cartilage rings (1). To assess whether hillocks are present in the young airway, we examined mouse trachea at postnatal day 3, 6, 14, and 28 (3). We then sought to identify islands of airway epithelium characterized by Krt13 protein expression in aciliate patches of epithelium defined by the absence of ciliated cells marked by acetylated tubulin staining (Fig 1B). Indeed, postnatal day 3 (P3) trachea contain aciliate Krt13+ patches of cells.

**Figure 1.**
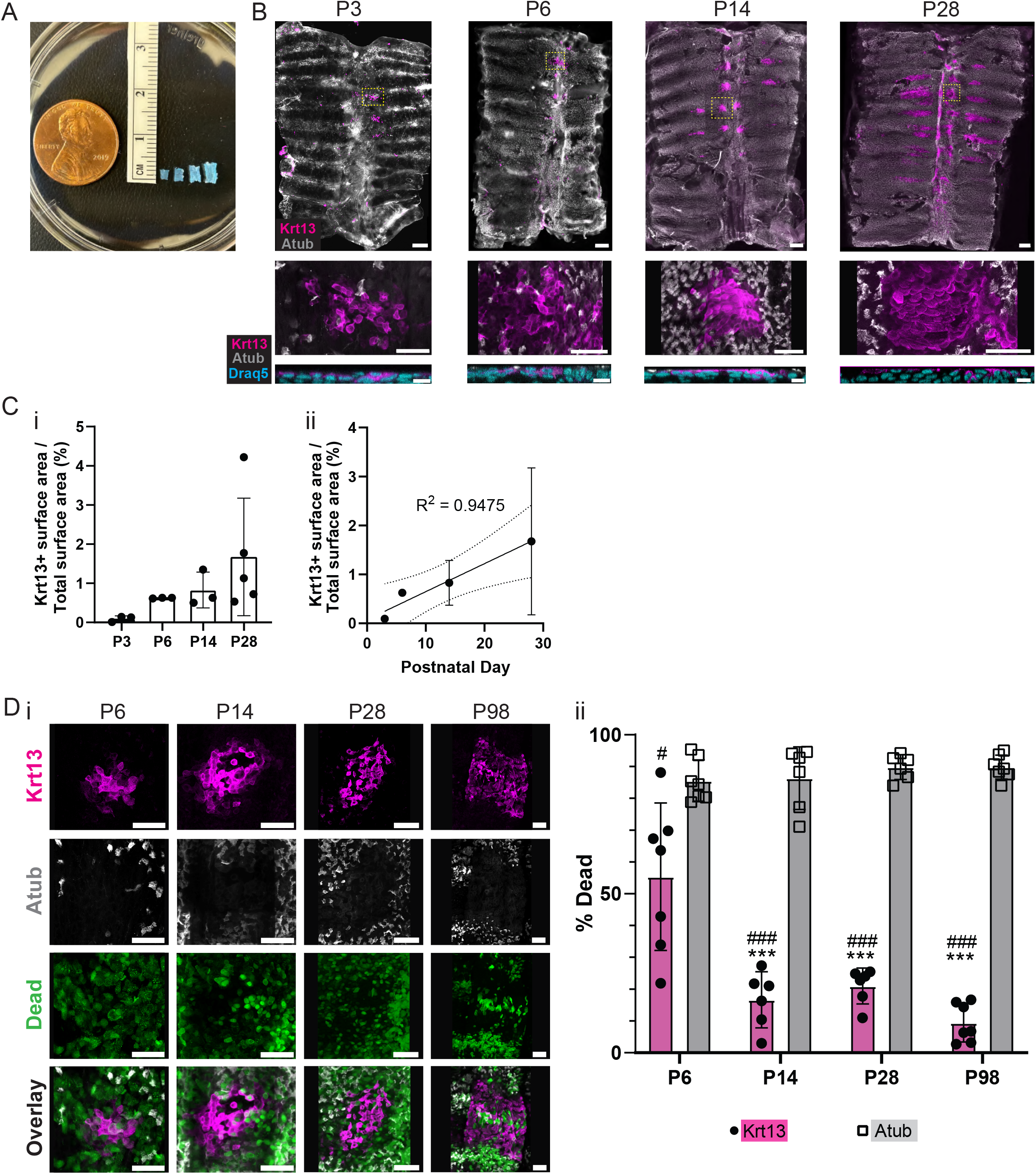
Postnatal development of murine hillocks. (A) Relative sizes of trachea wholemounts from P3, P6, P14, and P28 day old mice. (B) Top panels: Staining for keratin 13 (Krt13, magenta) and acetylated tubulin (atub, gray) in P3 to P28 murine tracheal wholemounts demonstrates the presence of hillocks in characteristic locations over and adjacent to posterior membrane and cartilage rings. Scale bar 200µm. Middle panels: Magnified images of individual hillocks (boxed in upper panels) at each time point. Scale bar 50µm. Lower panels: Optical cross sections demonstrating stratification of nuclei (draq5, cyan). Scale bar 10µm. (C) (i) Percentages of total surface area occupied by Krt13+ hillocks increase with time. (ii) Linear regression consistent with a linear increase in Krt13 surface area % over time. Dotted line represents 95% confidence interval. R^2^ = 0.945. (D) Hillocks acquire injury resistance postnatally. (i) Representative hillock whole mounts (Krt13, magenta) are injury resistant following acid injury and contain fewer dead cells (Live or Dye stain, green) than the surrounding ciliated epithelium (Atub, gray). (ii) Quantification of dead cells in hillocks and ciliated epithelium. Hillocks resist injury compared to surrounding ciliated cells at all ages (paired t-test; # p<0.05, ### p<0.001). Hillocks in P6 explants are less injury resistant than older explants (ANOVA, Tukey’s multiple comparison test; ***p<0.001) as seen in Di. N = 6-7 hillocks at each age.

High resolution cross-sectional imaging reveals that these Krt13+ patches were comprised of stratified cells with flattened nuclei mirroring their adult squamous counterparts (Fig 1B). The airway epithelial surface area covered by the Krt13+ cells increases linearly from P3 to P28 (Fig 1Ci-ii). Notably, the percent of the airway surface occupied by hillocks at P28 is similar to that seen in adult mice (1.27 – 4.46%) (1).

Since adult hillocks are characterized by their resistance to injury, we sought to assess the injury resistance of developing hillocks. As infants frequently experience aspiration following reflux, we employed an acid injury protocol in postnatal tracheal explants as we previously performed using adult airway (1). Tracheal explants were treated with 80mM HCl for 2 minutes, after which cell death was quantified in hillocks relative to surrounding ciliated epithelium (Fig 1Di-ii). The consistency of the acid injury was confirmed by measuring the percentage of dead ciliated cells. This percentage did not vary with age (85-90% at each timepoint) (Fig 1Dii). In contrast, hillocks were resistant to acid injury from P6 through P28. However, quantification of the percentage of dead cells showed that P6 hillocks were less injury resistant (55% of Krt13+ cells were dead) than older P14, P28 and P98 hillocks (9-20% of Krt13+ cells were dead). Thus, P6 hillocks are functionally immature.

We next sought to identify infant and pediatric human hillocks. Using a dataset of cells that previously annotated hillock precursor cells, hillock cells, cycling hillock cells, and squamous cells from donors across the human lifespan, we subset 15,711 epithelial cells that originated from tracheal and bronchial brushings of 11 healthy pediatric donors (1 month to 16 years old) and integrated the data to remove batch effects (4). We then performed dimensionality reduction and clustering, followed by uniform manifold approximation and projection (UMAP) to visualize the data in two-dimensional plots. Clusters were defined based upon the expression of canonical cell type markers. We identified KRT13+ airway epithelial cells at all ages. These cells were found within three clusters (Fig 2Ai-ii). One cluster was characterized by the expression of squamous cell markers and named hillock squamous. A second cluster displayed less enrichment of squamous cell markers and the coincident low level expression of secretory cell markers including SCGB1A1, and thus was named hillock transitional. The final population of KRT13+ cells mapped onto the basal cell cluster, representing hillock basal cells. Notably, some of these KRT13+ cells found within the basal cell cluster could represent the previously reported differentiating KRT13+ suprabasal intermediates of the pseudostratified epithelium, however, this cannot be resolved without additional spatial information (1, 5). We then defined a gene set comprised of the most highly expressed genes of the human hillock squamous cluster (Fig 1Aiii). A corresponding gene set of murine orthologs was projected onto a UMAP of mouse tracheal epithelium and mapped onto the murine hillock luminal cluster with remarkable precision. To confirm that bona fide hillocks do indeed exist in the pediatric airway, we obtained mainstem bronchial tissue from an 8-month-old and an 8-year-old. In both samples, KRT13 staining revealed hillocks with their characteristic location and gross morphology (Fig 2Bi-ii).

**Figure 2.**
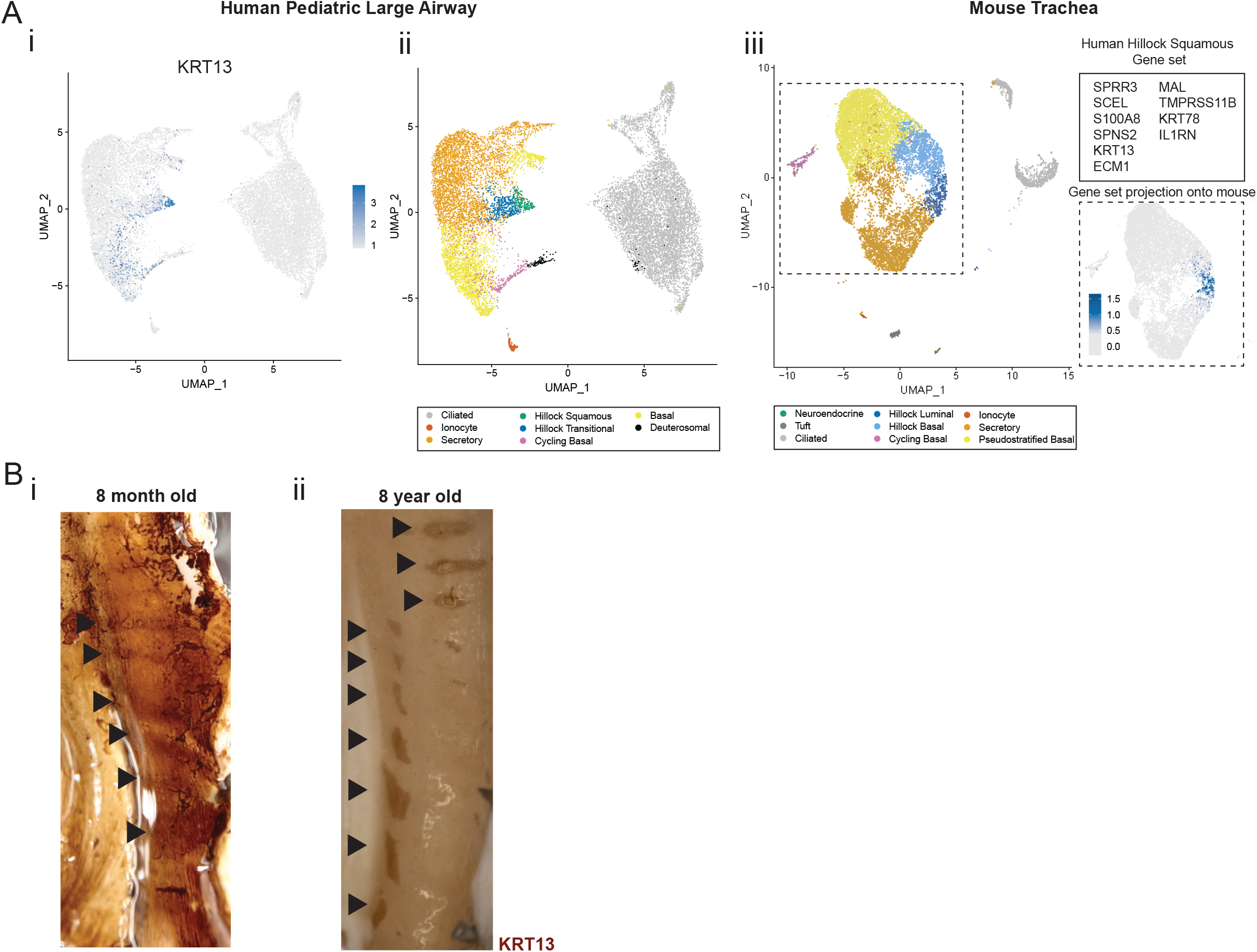
Hillocks are present in human pediatric donors from infancy through adolescence. A) Uniform Manifold Approximation and Projection (UMAP) plots of human (i, ii) and mouse (iii) airway epithelial cells. Human and mouse data are reanalyzed from previously published studies (1, 4). (i) Human airway epithelial cells from tracheal and bronchial brushings of healthy pediatric donors (1 month to 16 years) were assessed for expression of the conserved hillock marker KRT13. (ii) Human hillock squamous and hillock transitional cell clusters are identified. (iii) A gene set comprised of human hillock squamous cell markers was defined and a corresponding gene set of murine orthologs mapped directly onto murine hillock luminal cells (arrows). (B) (i) DAB (KRT13, brown) staining of wholemounts of 8-month-old and (ii) 8-year-old mainstem bronchi reveal characteristic KRT13+ hillocks (arrowheads).

Of note, KRT13+ cells have been described in pediatric and adult asthma, occurring as ill-defined clusters of cells without a classic hillock morphology (6, 7). It will be interesting to establish whether there is a relationship between hillocks, differentiating KRT13+ suprabasal cells of the pseudostratified epithelium, and these disease-associated clusters of cells. Future work using spatial transcriptomics may clarify the relationship, or lack thereof, of these morphologically distinct cell populations. Historically it has been thought that squamous cells are rare in the young airway. However, a catalogue of samples from 2170 pediatric autopsies documented squamous patches in 37 donors, perhaps representing the earliest evidence for pediatric hillocks (8). One wonders whether the postnatal airway is more susceptible to damage from infection or aspiration due to the immaturity of hillocks (9). Given that we have identified airway hillock cells across the lifespan, it will be interesting to see whether hillock numbers and size vary with age and whether they correlate with age-associated resilience in the face of injury.

## Contributions

YO: analysis, acquisition, interpretation, writing, editing

GS: analysis, acquisition, interpretation, writing, editing

QL: acquisition

JS: acquisition

LPH: acquisition, editing

VSS: conception, analysis, acquisition, interpretation, writing, editing, overall supervision

JR: conception, interpretation, writing, editing, overall supervision

## Supplementary Material

## Acknowledgements

The authors would like to thank New England Donor Services and the patients’ families who graciously donated their organs for science. We thank Sunghyun Kim for his input, Jim Haber (Brandeis University) for his suggestions for Yohana Otto’s Senior Honors Thesis, and the members of the Rajagopal and Chivukula labs for their input.

## Methods

### Mice

C57/B6 mice (JAX: 000664) at various ages were employed in all murine experiments.

Mouse dissection and wholemount staining: As previously described (1, 3), adult mice were euthanized as per IACUC protocols using CO2 and subsequent exsanguination through transection of renal artery. Mouse pups were euthanized by decapitation. Trachea were excised as previously described. Trachea were then transected along the ventral midline and splayed to expose the lumen. These explants were then fixed in 4% PFA for 10 minutes, permeabilized in PBS with 0.03% triton for 10 minutes, and placed in blocking buffer (5% BSA with 0.03% Triton in PBS) with primary antibody either for 1 hour at 37 deg or overnight at 4 deg. For immunohistochemistry, antibody recognizing Krt13 (abcam, ab92551) was diluted 1:300 and antibody recognizing Atub (Sigma, T-6793) was diluted 1:500. After two PBS washes, samples were incubated in secondary antibody for 1 hour at room temperature (donkey anti-rabbit Alexa 594 1:500 [Thermo Fisher Scientific A-21207]; donkey anti-rabbit Alexa 488 1:500 [Thermo Fisher Scientific A-21206]; donkey anti-mouse Alexa 488 1:500 [Thermo Fisher Scientific R37114]; donkey anti-mouse 405 1:500 [Jackson Immuno, 715-475-150]; draq5 1:1000 [Thermo Fisher Scientific 62251]. Samples were mounted with Southern Biotech Fluoromouont-G 0100-01.

### Acid Injury

Tracheal explants were incubated in 80mM HCl for 2 minutes with agitation as previously described (1). Samples were immediately placed into tubes with DMEM and Biotium Live-or-Dye Nucfix Red staining kit (#32010, 1:1000) for 10 minutes. Samples were then fixed and stained per manufacturer protocol.

Human whole mount staining: Deidentified donor lungs were received from New England Donor Services with Mass General Brigham IRB approval. Informed consent was obtained by New England Donor Services. Airways were dissected from lungs. A single incision was made along the ventral aspect of the airways from the trachea to the mainstem bronchi. Airways were then pinned to a soft silicone mat poured in a 15-cm petri plate. A dam of modeling clay was used to outline the airway structure in order to minimize antibody use. Tissue was then fixed with 4% PFA for 1 hour with gentle agitation.

After 3 washes in PBS, tissue was permeabilized in 0.1% PBST (tween) for 1 hour with gentle agitation. Samples were incubated in 1:1000 KRT13 antibody diluted in 1% BSA, 3% triton in PBS and incubated overnight (4 C). Sample were then washed 3x in PBS for a total of 1 hour. Horseradish peroxidase-anti-rabbit antibody (1:1000) was then incubated with the tissue in TNB block (0.1M Tris, pH7.5, 0.15M NaCl, 0.5% casein blocking reagent) for 1 hour at RT with gentle agitation. Then, tissue was incubated with 2 mg/ml DAB dissolved in 0.1M citrate-HCL at pH5.1 for 20 minutes at RT with gentle agitation. Tissue was then washed 5x in PBS for a total of 2 hours at RT. A Nikon D850 digital camera with a Nikon DX AF-S Nikkor 55-300mm f/4.5-5.6G ED lens, aperture set to f/5.6, ISO 3200, shutter speed 1/500 s, focal length 300mm was used to photograph whole mount airways.

8-month-old: male patient who passed away from bacterial meningitis with positive blood cultures for *Streptococcus pneumoniae*.

8-year-old: male patient who passed away of unclear etiology. Suspected meningitis with preliminary negative blood cultures.

Pediatric scRNAseq reanalysis and comparison to murine scRNAseq: Pre-analyzed human single cell RNA sequencing data published previously (4) were downloaded via the Human Cell Atlas portal: explore.data.humancellatlas.org/projects/1538d572-bcb7-426b-8d2c-84f3a7f87bb0. The mouse single cell RNA sequencing dataset was reported in (1). All analysis was performed in R using Seurat v.4.3.0.1. The original human dataset contained matched samples from peripheral blood and nasal, tracheal and bronchial brushings of healthy and COVID-19 infected donors across the entire human lifespan. For our analysis, we included: 1) only the epithelial cell compartment, 2) from samples collected by tracheal and bronchial brushings, 3) from healthy pediatric donors (ages 1 month to 16 years), 4) samples that included a minimum of 100 epithelial cells. This resulted in a set of 15,711 epithelial cells from 11 healthy pediatric donors. To remove batch effects prior to analysis, we normalized the data using the SCTransform() function, selected 3000 integration features via the SelectIntegrationFeatures() function, and integrated via the IntegrateData() function. Principal component analysis was performed on the integrated dataset. The number of significant dimensions was estimated using the ElbowPlot() function, which plots the standard deviations of principal components. We then performed clustering via the FindNeighbors() and FindClusters() functions, and then uniform manifold approximation and projection (UMAP) to visualize the data in two-dimensional plots. Cluster identity was assigned based on the expression of the following epithelial markers: KRT5, TP63 (basal cells), SCGB1A1, MUC5AC (secretory/goblet cells), MKI67 (cycling cells), FOXJ1 (ciliated cells), ASCL1 (neuroendocrine cells), DEUP1 and FOXN4 (deuterosomal), and FOXI1 (ionocytes). To assign an identity to the hillock squamous cluster, we performed differential gene expression using FindMarkers(). The top 36 differentially expressed genes were analyzed using Enrichr, and the top cell type category was found to be “Squamous epithelial cells in lung” based on the expression of 26 markers including the hillock marker KRT13 and the squamous cell-associated genes ECM1, SPRR1b, 2a, 2d, and 3. The hillock transitional cluster was assigned based on the co-expression of a set of 9 squamous epithelial markers identified using Enrichr and the secretory cell gene SCGB1A1.

We next defined a set of differentially expressed genes enriched in the human hillock squamous cell cluster. To do so, we extracted the top differentially expressed genes for each cluster using the FindMarkers() function and calculated a prevalence score for each gene according to the following formula (as described in (3)):

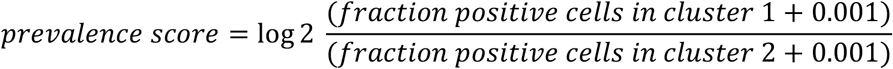

We then ranked the genes according to both the prevalence score and RNA expression levels. We selected the top differentially expressed genes to define a human hillock squamous cell gene set (see Figure 2A iii). The enrichment of the orthologous mouse gene set was assessed using our previously published murine dataset via the AddModuleScore() function. The module scores were then projected onto a UMAP of murine airway epithelia.

